# ENDOMETRIAL GLANDULAR EPITHELIAL CELL PLASTICITY DURING POSTPARTUM REGENERATION AND ESTROUS CYCLICITY

**DOI:** 10.64898/2026.10.07.757385

**Authors:** N Srivastava, D Sinzu-Prieto, CA Pru, TE Spencer, AM Kelleher, JK Pru

## Abstract

Several different mechanisms of regeneration have been proposed to explain repair of the endometrial luminal epithelium (LE) following menstrual/estrous cyclicity, parturition or other damaging insults to the uterine mucosa. These include activity of an as-yet unidentified epithelial stem cell population, mesenchymal transdifferentiation, or integration of bone marrow-derived cells. Here, we used a novel *Cag-Flox-STOP-Flox-Pgrmc2-3XFlag* (*Pgrmc2-3XFlag*) reporter mouse coupled with a recently developed GE-specific cre recombinase mouse (*Prss29-Cre*) to determine if and to what extent the GE contributes to regeneration of the LE following parturition and during aging in nulliparous mice. To initiate regeneration in the endometrium, control (*Pgrmc2-3XFlag*) and reporter (*Prss29-Cre;Pgrmc2-3XFlag*) mice were placed with males for one pregnancy. Postpartum (pp) uterine tissues were then collected 5 days (1x5dpp), 3-weeks (1x3wpp) and 7-weeks (1x7wpp) later. A fourth set of uterine tissues was collected 3-weeks after three pregnancies (3x3wpp). The number of glands did not change at any time point. Indelibly marked FLAG^+^ GE cells were detected by immunohistochemistry and immunofluorescence in GE as expected. As demonstrated in serial uterine sections of uteri from all four treatment groups, GE cells migrated toward the LE and formed junctional points (JP) where GE cells transitioned from cuboidal GE cells to pseudostratified columnar LE cells. Transitioning cells at junctional points expressed markers of both GE (*i.e.*, FOXA2) and LE (*i.e.*, CALB1) tissues. Following transition, GE-derived LE cells expressed PGR and ESR1 consistent with surrounding developmentally-derived LE cells. About 17 FLAG^+^ LE cell islands were also observed in 1x5dpp 4 mm longitudinal sections at sites distant from GE-LE junctional points. Interestingly, the number of islands did not change from 1x5dpp to 1x7wpp. However, the total number of GE-derived FLAG^+^ LE cells observed in each island decreased from 1x5dpp to 1x3wpp, but then tended (*p*=0.07) to increase by 1x7wpp. There was no difference in the number of GE-LE junctional points or GE-derived FLAG^+^ LE islands between 1x3wpp and 3x3wpp tissues. However, a significant increase in the total (*p*=0.02) and average (*p*=0.02) number of FLAG^+^ LE cells was observed in 3x3wpp LE islands highlighting the proliferative expansion of these cells over the intervening ∼80 days. GE-LE junctional points were also observed in 5-month old nulliparous *Prss29-Cre;Pgrmc2-3XFlag* mice at diestrus, but not in younger 6-week old mice. Collectively, these findings establish the GE as a quantifiable cellular source for LE regeneration during postpartum involution and as part of normal LE cell turnover during the estrous cycle in nulliparous mice. We propose a mechanism whereby GE cells within glandular structures are not terminally differentiated, but rather are stably differentiated to perform glandular function. However, when needed and based on the level of endometrial damage, GE cells can dedifferentiate, become integrated in to LE tissue and then reprogram to take on an LE phenotype as part of the regenerative process following parturition and during estrous cyclicity over time.

## INTRODUCTION

Throughout the female mammalian reproductive lifespan, the endometrium undergoes dramatic regenerative repair during the estrous/menstrual cycle, pregnancy, and following parturition [1, 2]. It is one of the few tissues that undergoes natural cycles of repair without significant scarring [3]. Post-menstrual repair in women is initiated almost immediately at the end of the cycle and is completed when bleeding ceases by around day five of the next cycle [4]. In contrast, and likely due to much more extensive damage, human postpartum regeneration begins immediately after delivery of the placenta and is completed over the course of several weeks [5]. While murine implantation is eccentric and human implantation is interstitial, both species engage in invasive hemochorial placentation where the trophectoderm actively invades the deciduum as it works to interdigitate with uterine vasculature to facilitate nutrient and waste exchange during fetal life [6]. In the mouse, the luminal epithelium (LE) is extruded by trophectoderm cell pseudopodia that physically dislodge LE cells from the basement membrane shortly after attachment [7, 8]. In humans, the trophectoderm actively secretes proteases to break down the basement membrane and displaced LE cells are then engulfed by the invading trophectoderm [9, 10].

At least one systemic and three local regenerative mechanisms have been proposed that describe how endometrial epithelia are reconstituted following natural cycles of damage [11–13]. First, a systemic mechanism suggests the activation and recruitment of stem or unspecified cells distantly housed in bone marrow to the uterus where they eventually differentiate into endometrial epithelial and stromal cells [14–18]. Second, a local stem cell-based mechanism proposes that mono-/bi-potent stem cells are activated following damage to expand, differentiate and re-epithelialize the GE and LE compartments [19–21]. Third, transdifferentiation has been proposed whereby subluminal stromal or perivascular mesenchymal cells switch fates and contribute to re-establishing the endometrial epithelia following damage. [1, 22–27]. Hysteroscopic studies of the human endometrium during menses suggest that after deposition of a fibrinous mesh over the denuded stroma, new luminal epithelial cells arise from the underlying stroma [28, 29]. The occurrence of mesenchymal-to-epithelial transdifferentiation is further supported by single cell transcriptomics revealing that the mid-secretory human endometrium harbors cells that express both mesenchymal and epithelial cell markers [30]. Single cell RNA sequencing and lineage tracing studies also support the concept that mesenchymal-to-epithelial transdifferentiation contributes to repair of the LE during menstruation [31]. Fourth, GE and LE cells individually have the capacity to dedifferentiate and then redifferentiate or reprogram into the alternative endometrial epithelial cell lineage. Recently, the LE was shown to reconstitute the ablated GE in adult nulliparous mice. Here, a genetic approach was used to ablate the GE with diphtheria toxin. Following confirmation of GE ablation, new LE-derived GE was observed within 5-10 days supporting the concept that LE cells retain some level of plasticity sufficient to allow their transition into GE following a damaging insult [32]. In opposition to LE-to-GE transition, Novak and Te Linde first described over a century ago that the LE could arise from the basalis GE [33]. Hartman later used surgical ablation of the non-human primate endometrium to demonstrate that deep residual basal GE stubs near the myometrial interface could reconstitute both the GE and LE in the overlying functionalis even after multiple surgeries [34]. Several more recent descriptive studies show remnant GE cells contributing to the re-epithelialization of the LE following menses in the human and non-human primate models [35–38]. Using a recently develop GE-specific *Cxcl15-iCre* mouse line in which approximately 86% of the GE cells express cre recombinase, the GE was shown to minimally contribute to regeneration of the LE following parturition; however, the extent was not quantified over time [39].

The current study was conducted to quantify GE contributions to the LE during postpartum endometrial regeneration, and to evaluate the extent of clonal expansion of these GE-derived cells over time. In order to accomplish this, a 3XFLAG-tagged reporter construct was expressed exclusively in the GE using the *Prss29-iCre* mouse, which indelibly labels 100% of GE cells during pregnancy [40]. It was also of interest to determine the extent of GE-to-LE transition over time in the aging nulliparous endometrium in order to determine if this transition is a component of normal LE turnover in the absence of significant tissue damage.

## MATERIALS and METHODS

### Animals

All animal experiments were approved by the University of Wyoming Institutional Animal Care and Use Committee. CD1 mice were purchased from Charles River Laboratories (Wilmington, MA) for evaluating uterine expression of *Prss29* mRNA. Uterine tissues were collected on postnatal (PND) 21, 2 months (mo), 6 mo, 12 mo, gestation day (GD) 7 and 3 weeks after primiparous parturition. Transgenic mice with a *Cag-loxP-STOP-loxP-Pgrmc2-3XFlag* construct inserted into the *Hip11* intergenic region were commercially generated at Applied StemCell using the TARGATT System. A PCR-based genotyping strategy was developed to identify control and transgenic mice. These FLAG reporter mice were crossed with recently developed *Prss29-Cre* mice to faithfully recombine the *Cag-loxP-STOP-loxP-Pgrmc2-3XFlag* construct and express PGRMC2-3XFLAG in the glandular epithelium (GE) [40]. Two or five month old control (*Prss29^+/+^;Cag-loxP-STOP-loxP-Pgrmc2-3XFlag)* and *Pgrmc2-3XFLAG* (*Prss29^Cre/+^;Cag-loxP-STOP-loxP-Pgrmc2-3XFlag*) female mice were placed with sexually mature and proven male CD1 mice to obtain pregnancy. Pregnant mice were allowed to deliver, and uterine tissues were then collected 5 days (1x5dpp), 3 weeks (1x3wpp) and 7 weeks (1x7wpp) after primiparous parturition or 3 weeks after a third pregnancy (3x3wpp) (see figure 2A). Tissues from all groups were collected during diestrus as determined from vaginal cytology and fixed for 24 h in 10% buffered formalin. Tissues were then temporarily stored at 4°C in 70% ethanol until embedded in paraffin wax using a HistoCore PEARL (Leica) tissue processor.

### *In Situ* Hybridization (RNAscope)

Formalin-fixed uterine tissues were embedded in paraffin wax. 5 µm thin sections were cut for in situ hybridization, using the Advanced Cell Diagnostics RNAscope 2.5 HD Reagent Kit-BROWN (Cat. #322300), according to the manufacturer’s instructions. Briefly, slides were deparaffinized in xylene and 100% ethanol. Hydrogen peroxide was applied for 10 minutes, followed by two H_2_O rinses. Antigen was retrieved using 1X Target Retrieval Buffer in a Decloaker (BioCare Medical) at 95°C for 15 minutes. Slides were air-dried, and sections were circled with hydrophobic pen. The RNAscope reagents Protease Plus, Mm-*Prss29*-C1 probe (Cat. #1110301-C1) and Hybridize AMP 1-6 were used as directed by applying and incubating in the HybEZ oven for specified times. DAB-A and DAB-B were mixed and applied to sections for 10 minutes at room temperature. Slides were placed in water to stop the reaction and then counterstained with Methyl Green for 20 minutes at room temperature. Sections were dehydrated in increasingly graded ethanol and xylenes before being mounted with Cytoseal 60 and coverslipped.

### Immunohistochemistry and Immunofluorescence

Paraffin embedded tissue sections (5 µm) were deparaffinized in xylenes and rehydrated in a series of decreasing ethanol washes and finally placed in phosphate buffered saline. Tissue sections were then subjected to antigen retrieval using a Decloaker pressure cooker at 95°C for 5 min in 10 mM sodium citrate. After cooling to room temperature, sections were quenched (30 minutes in 3% hydrogen peroxide and 97% methanol) and blocking reagent (10% bovine serum albumin, 0.1% normal goat serum, and 1% Triton X-100 in phosphate-buffered saline) was applied for 3 at room temperature. Sections were then incubated with primary antibody diluted in blocking solution overnight at 4°C in as outlined in **Table 1**. Slides were washed in 1x Tris-buffered saline with Tween-20 (TBST; 3 X 15 min) and incubated with secondary antibody. For immunohistochemistry, a biotinylated secondary antibody was applied for 30 min at room temperature. After washing as before, sections were incubated with the avidin-biotin-peroxidase complex (Vector Laboratories, CA) for 30 min and sections were then incubated in 3,3’-diaminobenzidine substrate (Vector Laboratories, CA). Sections were counterstained with hematoxylin, dehydrated in an ethanol gradient and xylenes, and mounted. For immunofluorescence, sections exposed to primary antibody were washed 3 X 15 min in TBST. A fluorescently tagged secondary anti-IgG immunoglobulin was applied under low light conditions followed again by 2 washes with TBST and one rinse with reverse osmosis water. Antibody specifications and dilutions are provided in Table 1. Slides were then mounted with Diamond anti-fade. For the anti-FLAG mouse antibody, blocking was performed according to the manufacturer’s instructions for the mouse-on-mouse (MOM) kit (Vector Labs, CA). Here, sections were incubated for 30 min in the primary antibody diluted in the MOM diluent. The secondary antibody incubation steps were the same as described above. Imaging of the slides subjected to immunohistochemistry was performed using an Axio Imager D2 (Zeiss) microscope at 20x and 40x magnification under brightfield illumination. The slides that underwent immunofluorescence labeling were imaged using an Olympus IXplore confocal microscope at 60X magnification. Images were combined using ImageJ (Fiji).

**Table 1.**
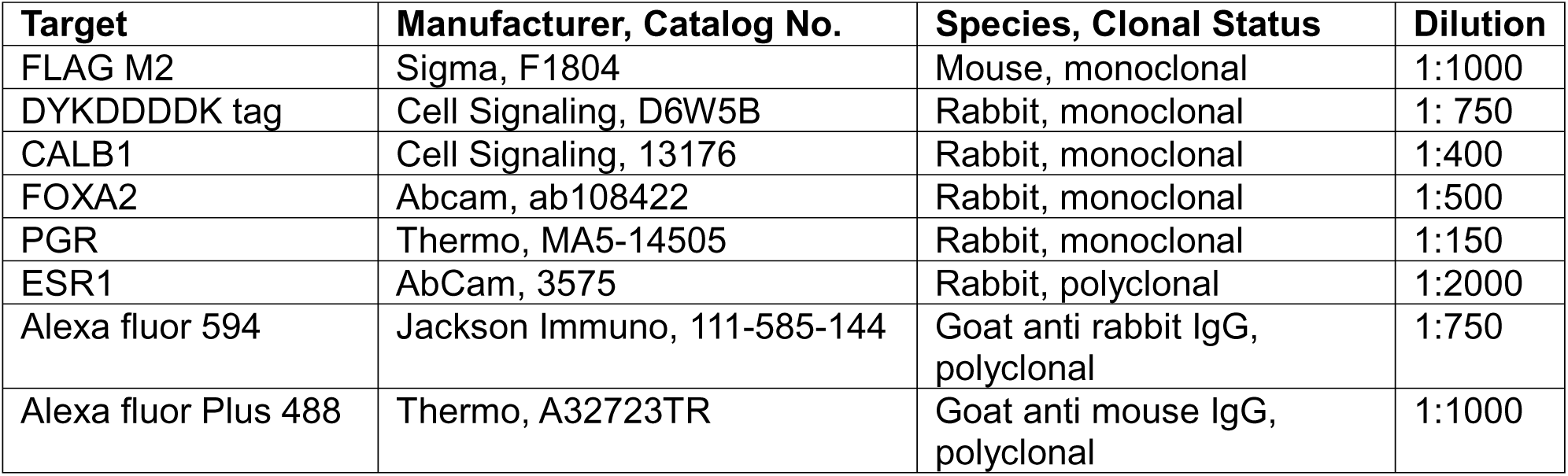
Antibodies.

Antibodies
| Target | Manufacturer, Catalog No. | Species, Clonal Status | Dilution |
| --- | --- | --- | --- |
| FLAG M2 | Sigma, F1804 | Mouse, monoclonal | 1:1000 |
| DYKDDDDK tag | Cell Signaling, D6W5B | Rabbit, monoclonal | 1: 750 |
| CALB1 | Cell Signaling, 13176 | Rabbit, monoclonal | 1:400 |
| FOXA2 | Abcam, ab108422 | Rabbit, monoclonal | 1:500 |
| PGR | Thermo, MA5-14505 | Rabbit, monoclonal | 1:150 |
| ESR1 | AbCam, 3575 | Rabbit, polyclonal | 1:2000 |
| Alexa fluor 594 | Jackson Immuno, 111-585-144 | Goat anti rabbit IgG, polyclonal | 1:750 |
| Alexa fluor Plus 488 | Thermo, A32723TR | Goat anti mouse IgG, polyclonal | 1:1000 |

### Quantification of GE-to-LE Transition during Endometrial Regeneration

For quantifying specified histological parameters such as the number of glands, junctional points, and luminal GE-derived cells in postpartum and aging nulliparous uterine sections, immunohistochemistry was first performed using the anti-DYKDDDDK (FLAG) antibody on 30 serially sectioned longitudinal uterine histological sections that were generated from a 4 mm section of the uterus. A total of 4 non-adjacent tissue sections separated by 9 intervening sections were used for quantification from n=3-4 replicates per group. Positively stained cells appeared brown after DAB staining. Junctional points (JP) were defined as the points where positively stained GE cells clearly contributed cells to the LE as determined by cellular morphology. LE islands were described as clusters of ≥2 positively stained pseudostratified LE cells. Both the number of LE islands and the number of cells within each island were counted. Because LE islands were sometimes observed at the JP, they were also included in the analyses. The average number of cells per LE island was calculated by dividing the total number of LE island cells by the number of LE islands. For quantifying the percentage of FLAG^+^ cells following parturition, LE nuclei were counted in three randomly selected, non-overlapping 40 µm LE segments per replicate (n=3; 9 total regions per group), and the mean LE nuclear density was estimated to be 0.24, 0.23, 0.20, and 0.26 nuclei/µm in the 1x5dpp, 1x3wpp, 1x7wpp, and 3x3wpp groups, respectively. The perimeter of LE was traced in each section using the freehand tool in Fiji. The total number of LE cells per section was calculated by multiplying the mean LE nuclear density (nuclei/µm) by the total LE length (µm). FLAG^+^ LE cells were manually counted. The percentage of FLAG^+^ LE cells per time point was calculated as: FLAG^+^ LE cells per time point (%) = (Avg # of FLAG^+^ LE cells/Avg # of total LE cells) x 100%. For assessing FOXA2^+^ LE cells following parturition, the total number of cells was estimated in two fields of view for each of 3-4 biological replicate in each group. FOXA2^+^ cells were manually counted as a percentage of total cells.

### Statistical Analyses

Animals with the same genotype were randomly assigned to the described treatment groups. All data are presented as the mean ± SEM for n=3–4 replicates in which individual animals represented a single biological replicate. A Student’s *t*-test assessed differences between the means of two groups, while comparisons of means of 3 or more groups were evaluated by a one-way analysis of variance (ANOVA) followed by Fisher LSD *post-hoc* test. A p-value of ≤ 0.05 was considered statistically significant regardless of the statistical test used. All statistical analyses were performed using GraphPad Prism 6.0 (San Diego, CA).

## RESULTS

### Development of a GE Lineage Tracing Model

We first generated a transgenic mouse line to indelibly mark and trace the fate of the endometrial GE lineage in adult mice following parturition and over time in nulliparous animals. Here, the C-terminus of the endoplasmic reticular protein PGRMC2 was tagged with three tandem copies of the FLAG sequence (PGRMC2-3XFLAG). The *Cag-loxP-STOP-LoxP-Pgrmc2-3XFlag* construct was targeted to the *Hip11* intergenic region (**Figure 1A**). Mice harboring the *Cag-loxP-STOP-LoxP-Pgrmc2-3XFlag* construct were crossed with *Prss29-Cre* mice to direct recombination at *loxP* sites and removal of the transcriptional STOP sequence exclusively in the endometrial GE in gravid female mice. Resulting *Prss29-Cre;Cag-loxP-STOP-LoxP-Pgrmc2-3XFlag* (*Pgrmc2-3XFlag*) mice were completely fertile [9.46 ± 0.68 SEM pups/litter versus 8.06 ± 1.17 SEM pups/litter (*p* = 0.33) for control and *Pgrmc2-3XFlag* mice, respectively] with no observable health issues indicating that long-term expression of the fusion protein was well tolerated. We have also crossed *Cag-loxP-STOP-LoxP-Pgrmc2-3XFlag* mice with *Pgr-Cre* mice [41], which express PGRMC2-3XFLAG in the female reproductive tract, mammary epithelium and gonadotrophs, and *Hprt-iCre* [42], which allows for global expression of PGRMC2-3XFLAG (Pru, unpublished). These two additional groups of mice were comparably fertile with respective control mice and again did not show any observable health issues throughout their lifetime. Uteri from all three groups of recombined mice were histologically normal compared with uteri from corresponding control mice.

**Figure 1.**
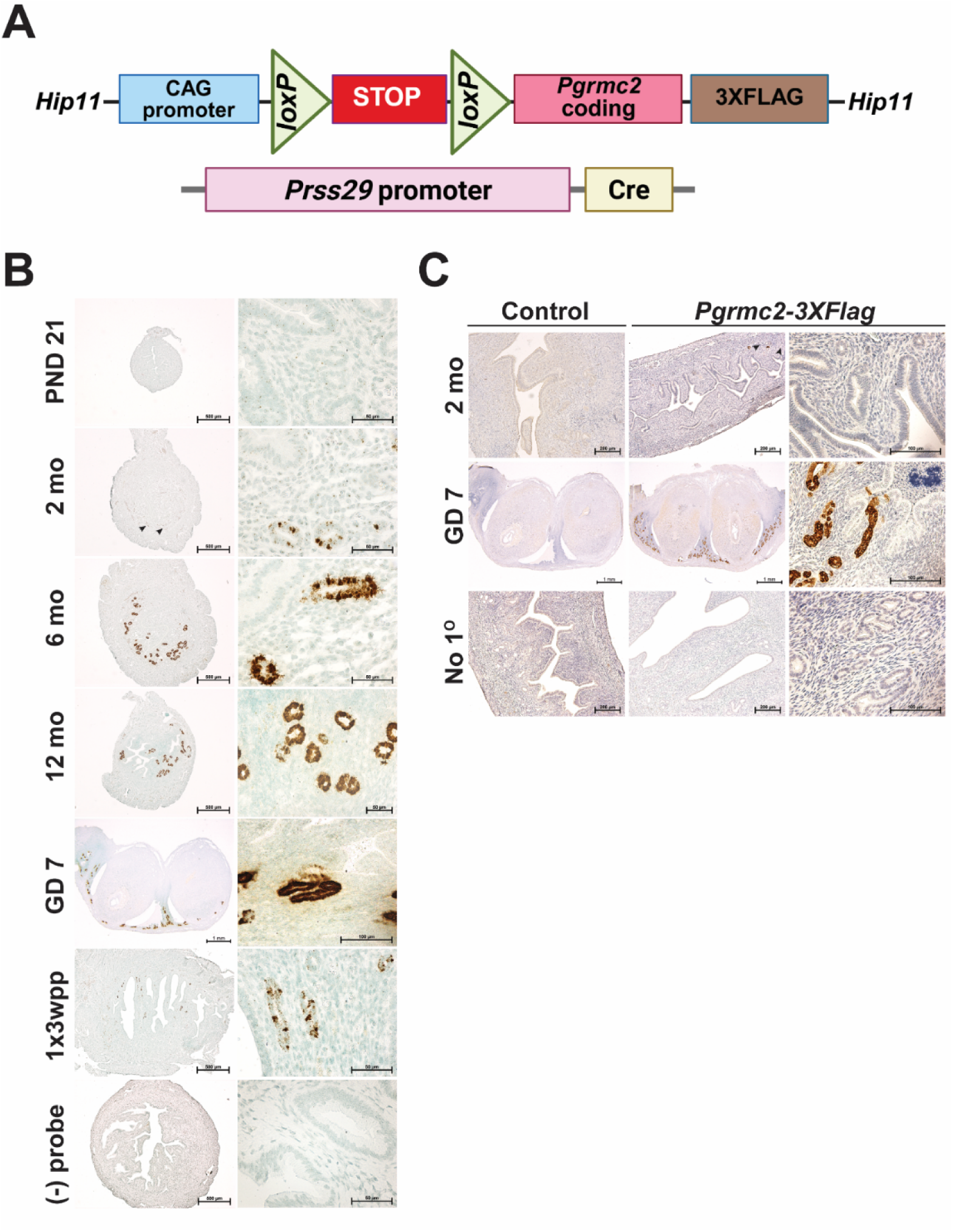
An endometrial GE lineage tracing model. (**A**) Mice harboring the *Cag-loxP-STOP-LoxP-Pgrmc2-3XFlag* construct were crossed with *Prss29-Cre* mice to direct recombination at *loxP* sites and removal of the STOP sequence exclusively in the endometrial GE in gravid female mice. (**B**) In the resulting *Pgrmc2-3XFlag* mice, RNAscope was used to detect *Pgrmc2-3XFlag* mRNA in GE tissue on postnatal (PND) 21, 2 months (mo), 6 mo, 12 mo, gestation day (GD) 7 and three weeks post-parturition of primiparity (1x3wpp). (**C**) Immunohistochemistry using anti-FLAG antibodies was used to detect PGRMC2-3XFLAG protein in uterine tissues obtained from 2 month old nulliparous and GD 7 primiparous control and *Pgrmc2-3XFlag* mice. Negative control results are also shown in which primary antibody was omitted (no 1). Scale bars represent 500 µm and 50 µm (**B**), 200 µm, 100 µm and 1 mm (**C**).

We first evaluated uterine expression of *Prss29* mRNA in nulliparous CD1 mice at PND 21 and 2, 6 and 12 months of age, as well as gestation day (GD) 7 and three weeks after primiparous pregnancy using RNAscope (**Figure 1B**). The *Prss29* mRNA expression was absent on PND 21 but then became increasingly expressed from 2 months to 12 months of age exclusively in GE cells and was absent in the LE, stroma and myometrium. Among the uterine tissues selected for analyses, *Prss29* mRNA was most abundant in the GE on GD 7. By three weeks post-partum (1x3wpp), *Prss29* mRNA expression was greatly diminished both in the level of expression on a per/cell basis, as well as the number of positively stained GE cells (**Figure 1B**). No signal was detected in *Pgrmc2-3XFlag* uterine tissue incubated without probe. As shown in **Figure 1C**, PGRMC2-3XFLAG protein was not detected by anti-FLAG IHC in uteri from nulliparous control or *Pgrmc2-3XFlag* mice at 2 months of age. However, PGRMC2-3XFLAG was abundantly detected in the GE of GD 7 primiparous mice at 2 months of age in *Pgrmc2-3XFlag* mice. The PGRMC2-3XFLAG protein was not detected in control mice that harbored the *Cag-loxP-STOP-LoxP-Pgrmc2-3XFlag* construct but lacked *Prss29*-driven cre recombinase expression or in uterine sections in which primary anti-FLAG antibody was omitted. These data demonstrate faithful *Prss29-*driven cre recombinase activity in the GE early in the first pregnancy.

### GE-to-LE Transition during Postpartum Endometrial Regeneration

Four different postpartum time points were evaluated for GE contributions to the LE and these included 5 days (1x5dpp), 3 weeks (1x3wpp) and 7 weeks (1x7wpp) after primiparous pregnancies, as well as 3 weeks after a third pregnancy (3x3wpp) (**Figure 2A**). In all cases, FLAG^+^ glands could be seen in serial longitudinal sections beginning in the peripheral stroma with subsequent sections showing fusion of these glands with and integration into the LE compartment (**Figure 2B-E**). It is interesting to note that the GE-to-LE transition continues to occur even 7 weeks after parturition, equivalent to about 11 estrous cycles, (**Figure 2D and 2E**). In some cases, the GE was found to extend beyond the LE into the luminal space at junctional points as shown schematically in **Figure 3A**. While there was no difference in the number of junctional points or LE islands derived from GE cells at 5 days, 3 weeks and 7 weeks beyond pregnancy, there was a significant decrease in the total number of GE-derived LE island cells and number of cells/island from 5 days postpartum to 3 weeks postpartum. Despite clear observation of glands contributing to the GE, there was no change in the number of glands within stromal tissue (**Figure 3B**). Interestingly, when comparing 1x3wpp and 3x3wpp uterine tissues, there was no difference in the number of junctional points despite much more long-term repair required after three pregnancies. Consistent with this finding, we also did not observe a change in the number of GE-derived islands within the LE compartment (**Figure 3C**). However, the total number of cells incorporated in GE-derived LE islands was increased about 4-fold in mice after three pregnancies compared with those having only a single pregnancy, presumably through clonal expansion. There was a corresponding 4-fold increase in the number of GE-derived cells within each LE island in mice with three pregnancies compared with single pregnancy female mice. The percentage of FLAG^+^ cells in the LE at 1x5dpp, 1x3wpp, 1x7wpp and 3x3wpp was 2.95%, 0.72%, 6.16% and 3.29%, respectively (**Figure 3B** and **3C**).

**Figure 2.**
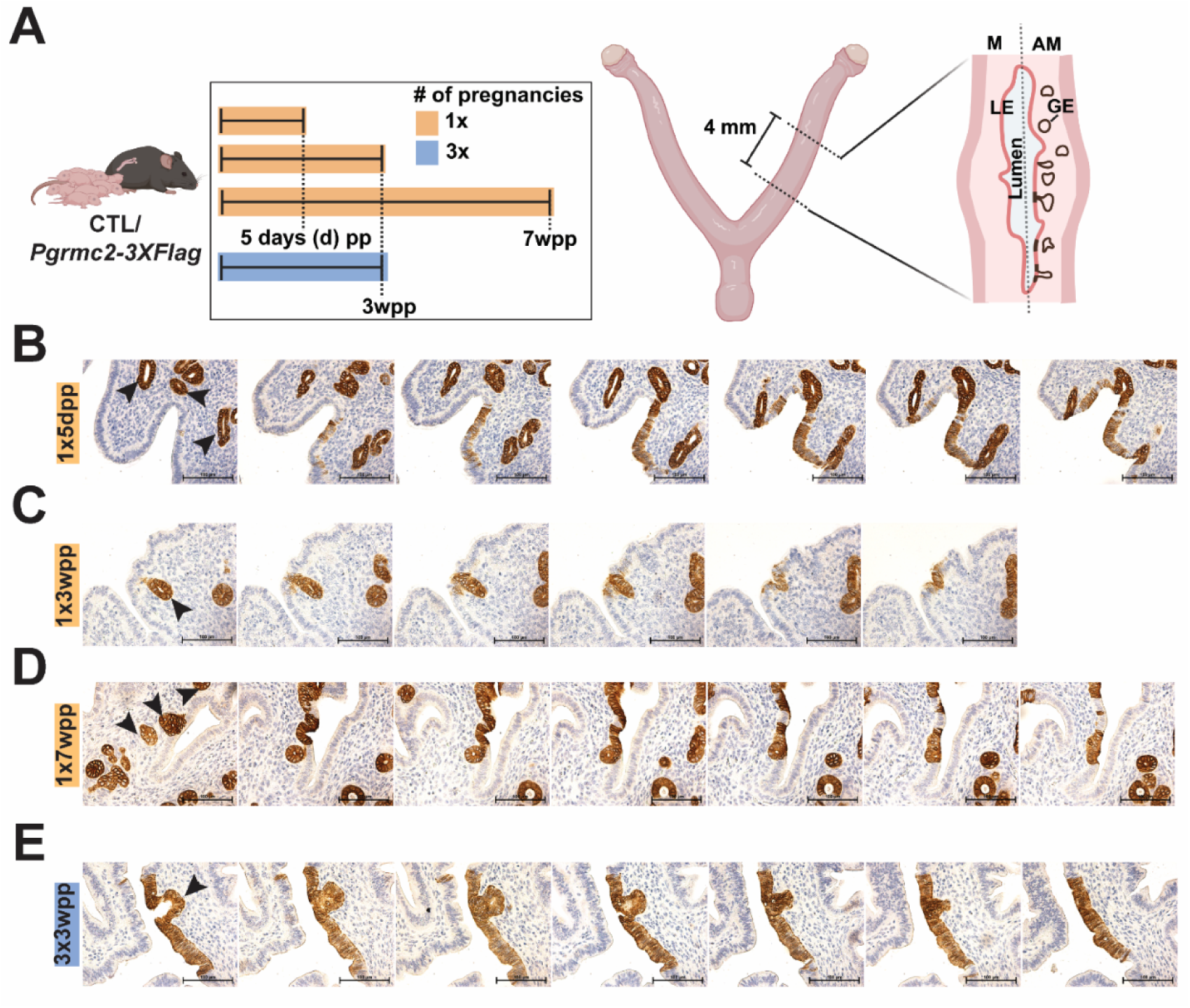
Lineage tracing of GE cells during postpartum endometrial regeneration. (**A**) Schematic showing timeline of when postpartum uterine tissues were collected from four groups of control and *Pgrmc2-3XFlag* mice following one or three pregnancies. A 4 mm section of uterus was obtained from each mouse and serial sectioned to assess contributions of the GE to the LE during regeneration using anti-FLAG immunohistochemistry. PGRMC2-3XFLAG protein detected in GE and LE in uterine serial sections obtained from *Pgrmc2-3XFlag* mice 5 days post-partum of first pregnancy (1x5dpp, **B**), 3 weeks post-partum of first pregnancy (1x3wpp, **C**), 7 weeks post-partum of first pregnancy (1x7wpp, **D**) and three weeks post-partum of third pregnancy (3x3wpp, **E**). Black arrows identify peripheral glands that fuse with LE. All scale bars represent 100 µm.

**Figure 3.**
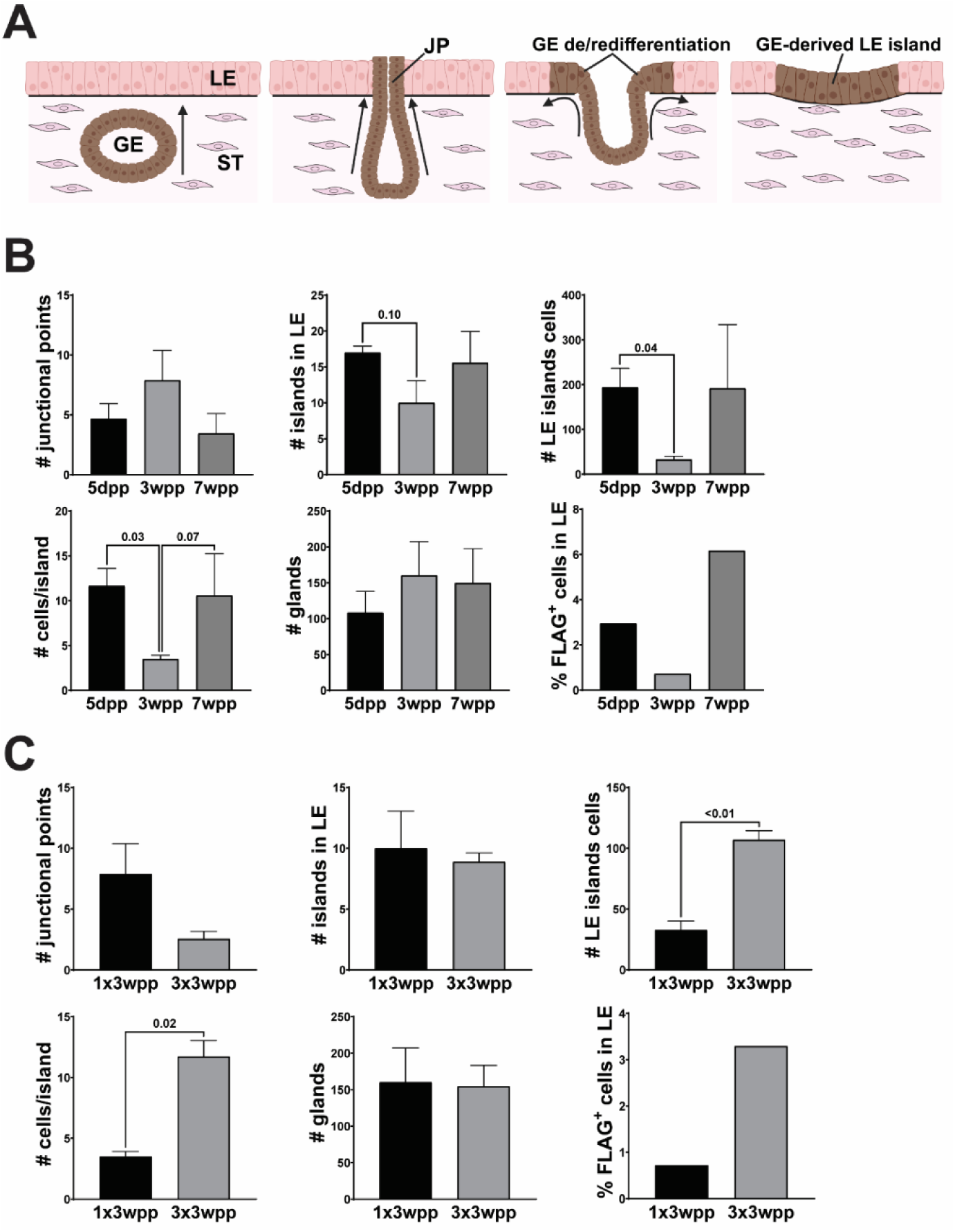
Quantification of GE-LE junctional points and GE-derived LE islands following parturition-induced endometrial regeneration. (**A**) Schematic showing transition of glandular epithelial (GE) cells to luminal epithelial (LE) cells following parturition. (**B**) Quantification of the number of junctional points, number of islands, number of GE-derived LE cells/island, number cells/island, number of glands counted and the percentage of FLAG^+^ cells in the LE in four non-adjacent uterine sections from each of 3-4 *Pgrmc2-3XFlag* mice 5 days, 3 weeks and 7 weeks after primiparous parturition. (**C**) Quantification of the number of junctional points, number of islands, number of GE-derived LE cells/island, number cells/island, number of glands counted and the percentage of FLAG^+^ cells in the LE in four non-adjacent uterine sections from each of 3-4 *Pgrmc2-3XFlag* mice at 3 weeks after primiparous parturition (1x3wpp) and 3 weeks after a third pregnancy (3x3wpp).

### GE-derived LE Displays an Expression Pattern Consistent with Developmentally Derived LE

Progesterone receptor (PGR) and estrogen receptor alpha (ESR1) expression in the uterus is highly dynamic and compartmentalized [43, 44]. Confocal immunofluorescence (IF) microscopy revealed that PGR is abundantly expressed in stromal cells in control and *Pgrmc2-3XFlag* uteri collected 5 days after parturition (1x5dpp) with reduced expression in the GE and no expression in the LE (**Figure 4A**). Dual IF staining for FLAG and PGR in *Pgrmc2-3XFlag* mice shows that while GE cells in distant glands express modest levels of PGR, this expression became greatly reduced or lost as cuboidal GE cells moved near the LE (**Figure 4A**). The expression of PGR is then lost as cuboidal GE cells transition into pseudostratifed LE cells. Importantly, a similar GE-to-LE transition was observed in uteri obtained from control mice. Conversely, ESR1 expression remained high in the LE, GE, and stroma during 5 days after parturition. Dual IF in 1x5dpp uterine sections revealed constant ESR1 expression in the FLAG^+^ GE cells transitioning into LE (**Figure 4B**). A similar ESR1 expression pattern was observed during the GE-to-LE contributions at junctional points in control tissues.

**Figure 4.**
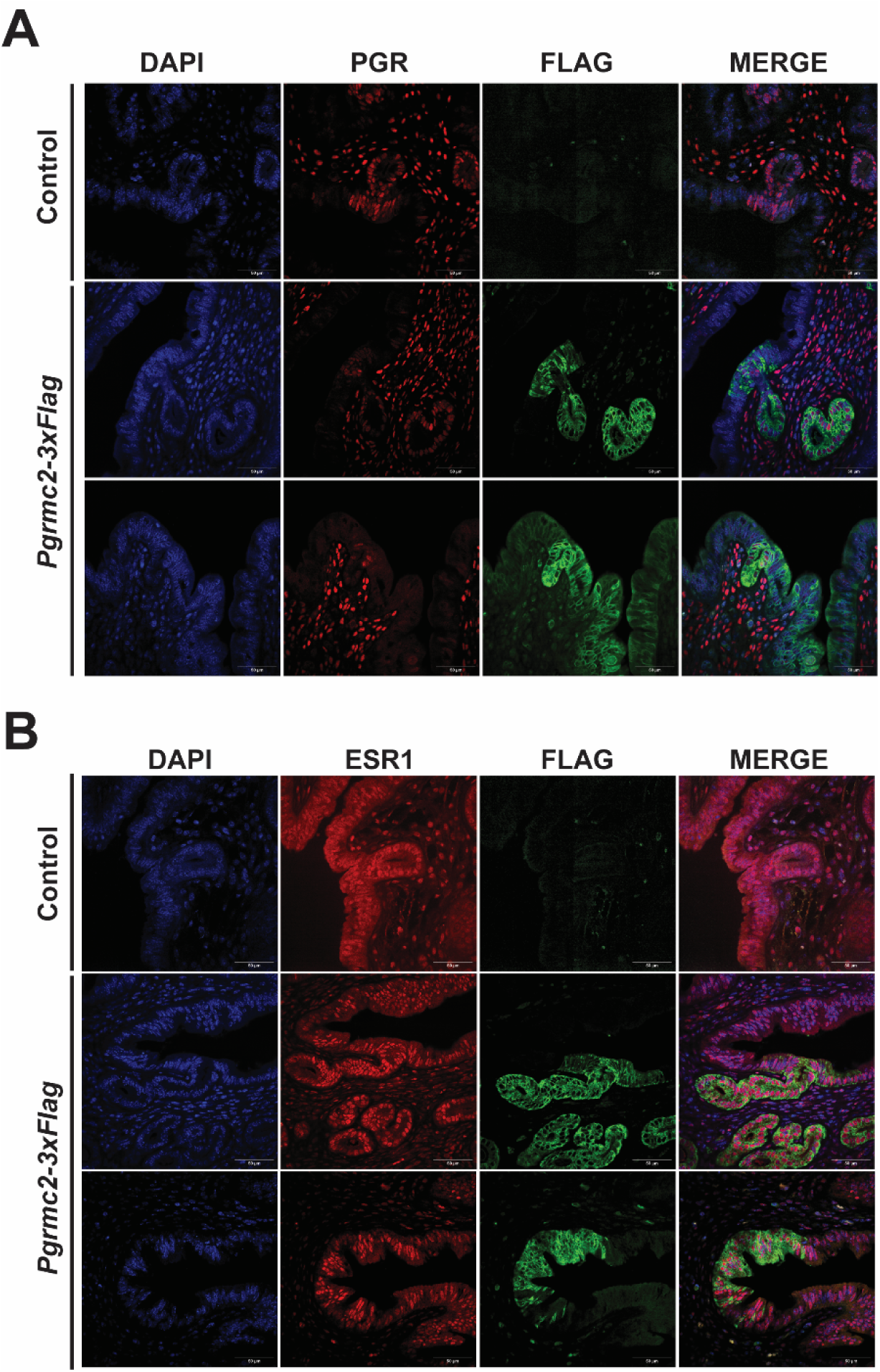
PGR and ESR1 expression in GE-to-LE transitioning cells. Immunofluorescent microscopy was used to evaluate expression of progesterone receptor (PGR) and estrogen receptor (ESR1) in control and *Pgrmc2-3XFlag* uterine sections obtained 5 days after primiparous pregnancy (1x5pp). Sections were co-stained with anti-FLAG antibody to identify GE-derived cells. n=3. All scale bars represent 50 µm.

Forkhead box A2 (FOXA2) is a well-established and widely used marker of endometrial uterine glands and regulates their development [45]. It was hypothesized that FOXA2 expression would gradually decline in the cuboidal GE cells undergoing GE-to-LE transition at junctional points. Dual IF for FOXA2 and FLAG in control and *Pgrmc2-3XFlag* uterine sections indeed revealed that GE cells transitioning to LE lost FOXA2 expression at junctional points 5 days after parturition (**Figure 5A**). In contrast, CALB1 is a marker of the LE with little to no expression in the GE [46]. As shown in **Figure 5B**, CALB1 was abundantly expressed in the LE of control and *Pgrmc2-3XFlag* uterine sections. In *Pgrmc2-3XFlag* sections, CALB1 expression was low to absent in FLAG^+^ GE cells distant from the LE, but then became abundantly expressed in FLAG^+^ GE cells as they transitioned from cuboidal GE cells to pseudostratified LE cells.

**Figure 5.**
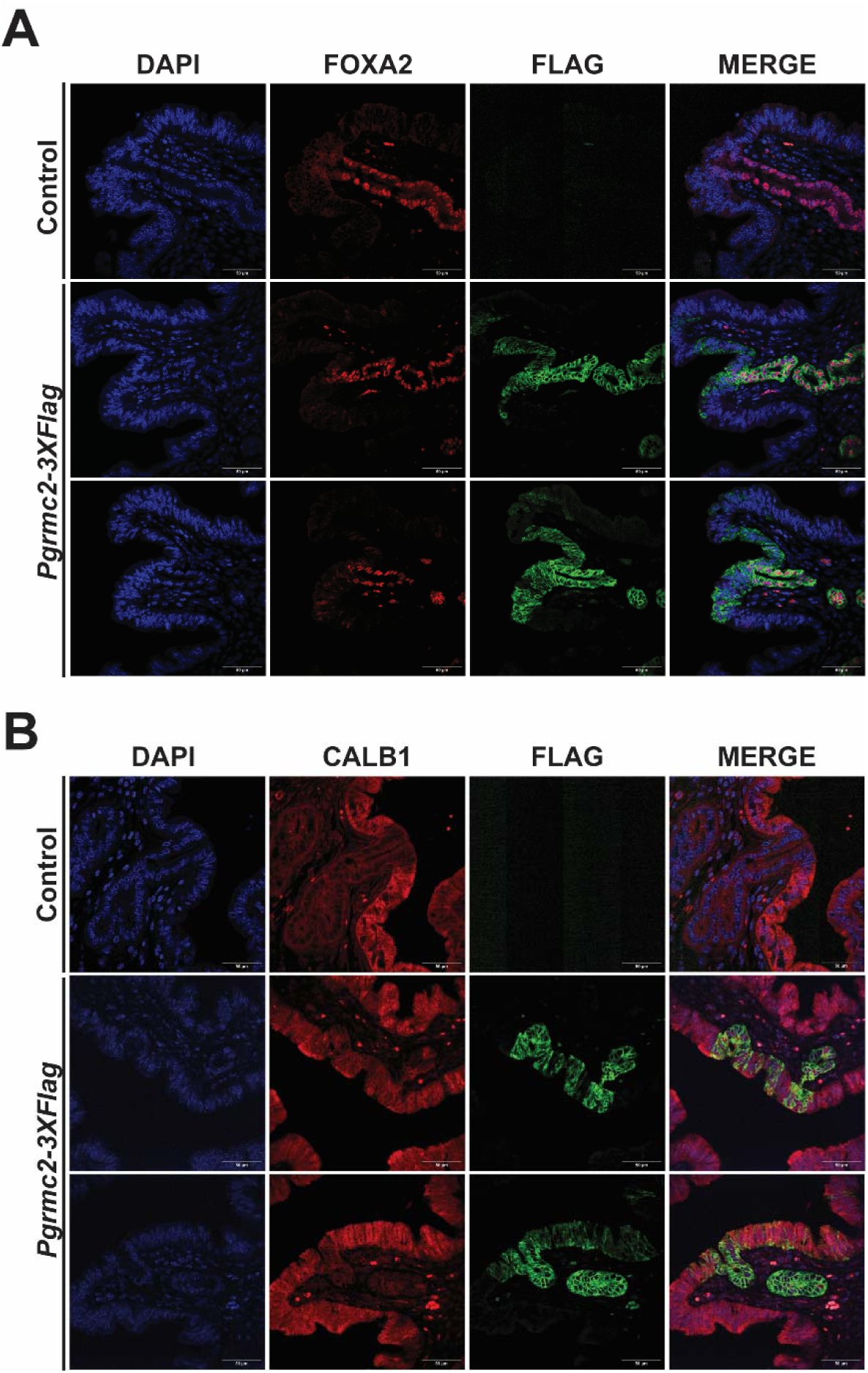
FOXA2 and CALB1 expression in GE-to-LE transitioning cells. Immunofluorescent microscopy was used to evaluate the expression of the GE marker forkhead box protein A2 (FOXA2) (**A**) and the LE marker calbindin-1 (CALB1) (**B**) in control and *Pgrmc2-3XFlag* uterine sections obtained 5 days after primiparous pregnancy (1x5pp). Sections were co-stained with anti-FLAG antibody to identify GE-derived cells. n=3. All scale bars represent 50 µm.

### The GE Traces to the LE with Age in Nulliparous Mice

While endometrial regeneration does not occur in estrous cycling animals as extensively as it does in menstrual cycling species or to the same extent as following parturition, there is still considerable turnover of the LE in mice through successive estrous cycles. We observed extensive PGRMC2-3XFLAG expression in nulliparous *Pgrmc2-3XFlag*, but not control, mice at five months of age indicating that *Prss29* cre recombinase activity is not completely restricted to pregnancy (**Figure 6A**). In *Pgrmc2-3XFlag* sections, FLAG^+^ GE glands can be seen in serial sections conjoining with and contributing cells to the LE at junctional points (**Figure 6B**). The GE-to-LE transition was age-dependent in nulliparous *Pgrmc2-3XFlag* mice. As shown in **Figure 6C**, very few FLAG^+^ GE cells were observed in the LE at 2 mo of age, suggesting that very limited GE-to-LE transition was occurring. However, the number of junctional points, GE-derived LE islands, total LE island cells, cells per island and FLAG+ glands all increased by 5 mo of age in nulliparous mice. In 4 mm longitudinal uterine sections obtained from 5 mo old *Pgrmc2-3XFlag* mice, we observed approximately 4 junctional points, 7 GE-derived FLAG^+^ LE islands, 7 FLAG^+^ cells/island and 95 FLAG^+^ glands.

**Figure 6.**
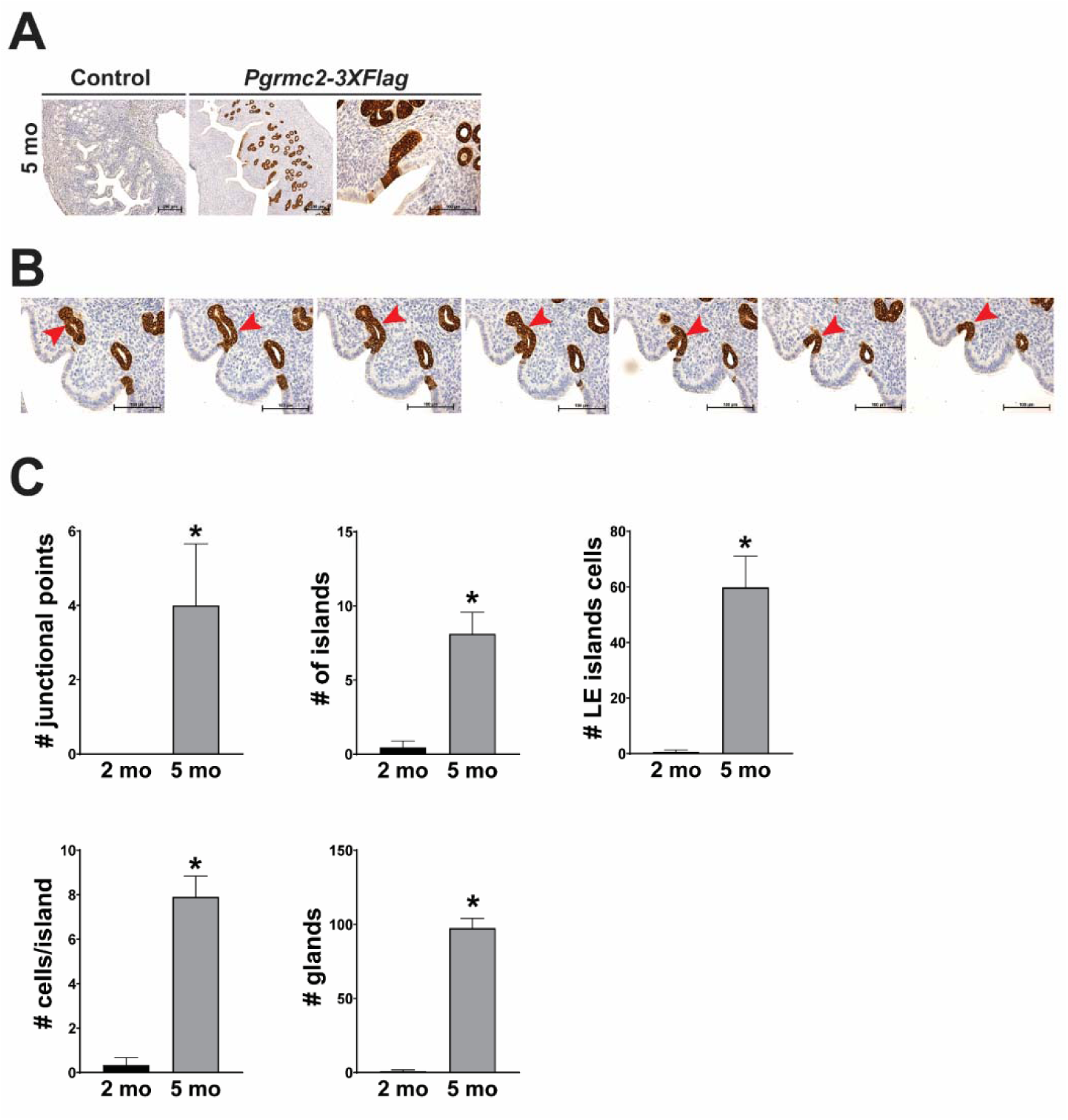
GE contributions to LE regeneration in nulliparous mice during aging. (**A**) Use of IHC to identify FLAG^+^ GE-derived cells in nulliparous control and *Pgrmc2-3XFlag* mice at 5 months (mo) of age. (**B**) Example of serial sectioned uterus showing progressive transition of GE into the LE of 5 mo old *Pgrmc2-3XFlag* mouse (red arrows). (**C**) Quantification of the number of junctional points, GE-derived LE islands, LE island cells, cells per island, and FLAG^+^ glands in 2 and 5 mo old nulliparous Pgrmc2-3XFlag mice. Data are representative of counts made from four non-adjacent (i.e., every tenth) serially cut sections. n=3. Scale bars represent 200 µm and 100 µm (**A**), 100 µm (**B**).

### Endometrial Regeneration Triggers FOXA2 Expression in the LE

Through the course of these studies, the interesting observation was made that FOXA2 was not exclusively restricted to the GE compartment, but rather became expressed in the LE 5 days after primiparous parturition (**Figure 7A and 7B**). Furthermore, FOXA2 expression in the LE was maintained as far out as 3 and 7 weeks beyond parturition (**Figure 7C and 7D**). The range of FOXA2^+^ cells in the LE varied within and between time points. FOXA2^+^ cells in uterine tissue obtained from control mice at 1x5dpp was between 2.89% and 23.92%. For *Pgrmc2-3XFlag* mice 3 weeks after primiparous parturition FOXA2^+^ cells ranged from 0.7% to 6.9% and at 7 weeks after primiparous parturition 1.16%-29.44%. We observed no difference in the number of junctional points or total glands in control and *Pgrmc2-3XFlag* mice 5 days after primiparous parturition (**Figure 7E and 7F**).

**Figure 7.**
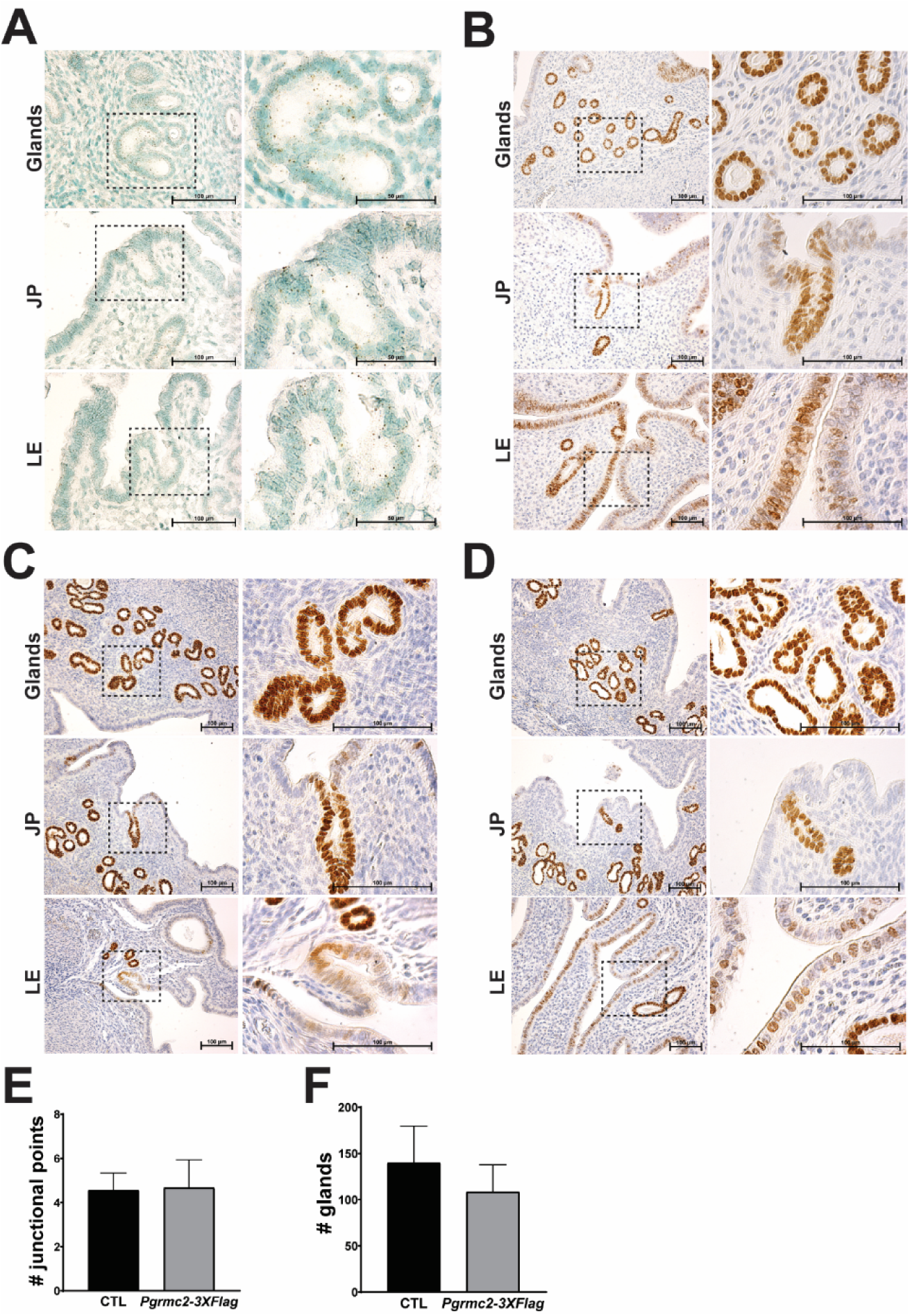
Endometrial regeneration triggers expression of the GE marker FOXA2 in the LE. RNAscope and immunohistochemistry were used to evaluate the expression of *FoxA2* mRNA (**A**) and FOXA2 protein (**B**) in glands, junctional points (JP) and LE in *Pgrmc2-3XFlag* uterine sections obtained 5 days after primiparous parturition (1x5dpp). Persistence of FOXA2 expression in endometrial GE, JP and LE after 3 (**C**) and 7 weeks (**D**) in primiparous post-parturient endometrium of *Pgrmc2-3XFlag* mice. Comparison of the number of GE-LE junctional points (**E**) and glands (**F**) between control (CTL) and *Pgrmc2-3XFlag* mice 5 days after primiparous parturition. All scale bars represent 100 µm.

## DISCUSSION

Failure to properly repair the endometrium has long-term negative reproductive consequences and reduces the quality of life in the female. For example, Asherman’s syndrome can occur following trauma or damage to the basal layer of the endometrium if LE tissue regeneration is compromised resulting in the development of intrauterine adhesions and excessive scarring in women [47, 48]. Negative energy balance in dairy cattle facilitates intrauterine bacterial infection following parturition which often results in endometritis and/or metritis, again caused by failure to quickly repair the LE barrier [49, 50]. In both cases, subsequent pregnancies become compromised because of the hyperinflammatory environment and long-term fibrotic damage to the tissue that generally does not resolve.

Several historical [33, 34, 51, 52] and more recent [20, 39, 53] studies have demonstrated that the GE participates in regeneration of the overlying LE when the endometrium has been naturally or artificially damaged by directly contributing glandular cells. The present study supports these prior findings and adds new data quantifying the extent to which the GE contributes cellularly to LE regeneration following single and multiple pregnancies, as well as in nulliparous mice where tissue turnover is more limited during the estrous cycle. A recent lineage tracing study using the GE-specific *Cxcl15-iCre* mouse demonstrated that the GE was capable of regenerating the entire LE following chemical ablation with polidocanol [39, 54]. However, GE contributions to the LE following during the estrous cycle (no more than 1.38%) and after artificial decidualization (0.27%) and subsequent involution were found to be minimal with limited clonal expansion once established in the LE. This is in contrast to the present study which suggests GE-to-LE transition occurs more frequently and that GE-derived LE cells clonally expand substantially over time. The average percentage of GE-derived LE cells (*i.e.*, FLAG^+^ cells) observed at 1x5dpp, 1x3wpp, 1x7wpp and 3x3wpp was 2.95%, 0.72%, 6.16% and 3.29%, respectively. Importantly, data presented in this study indicate that the GE contributes cellularly to the LE even in the absence of substantial endometrial damage as part of normal LE tissue turnover during the estrous cycle. As shown in **Figure 6**, while GE-to-LE transition was mostly absent at 2 months of age, this increased substantially by 5 months of age in nulliparous reporter mice. The discrepancy between the two studies may lie in the fact that while 100% of GE cells in the *Prss29-iCre;Pgrmc2-3XFlag* mouse used in the present study express reporter protein, only 86% of the GE cell in the *Cxcl15-iCre;R26^nTnG^*mouse express the reporter protein. It is possible that the remaining 14% reporter (*i.e.*, GFP) negative fraction of GE cells in *Cxcl15-iCre;R26^nTnG^* mice contribute significantly to the regenerating LE. While GE-to-LE transition does occur as part of normal cellular turnover during the estrous cycle, it does so at a reduced level compared to postpartum endometrial regeneration where much more tissue remodeling occurs. For example, the number of junctional points, GE-derived islands in the LE compartment, GE-derived cells, and GE-derived cells/island were 20%, 53%, 75% and 37% lower, respectively, in 5 mo old nulliparous mice compared with mice 3 weeks after primiparous parturition (1x3wPP). In the more extreme case described above, the GE has the capacity to regenerate the entire LE when chemically ablated [39]. Based on these collective findings, the level of GE-to-LE transition is evidently consistent with the amount of tissue damage. The symmorphotic factors and biophysical forces that allometrically pair the level of GE-to-LE transition with the amount of endometrial tissue damage remain to be determined, but it is likely that much of the instruction comes intrinsically from the epithelial cells and their position in the endometrium, as well as the mesenchymal compartment.

GE-derived cells at the junctional points retained intermediate GE-like PGR expression and sustained ESR1 expression as they transitioned into the LE (**Figure 4**). This was coupled with progressive downregulation of FOXA2 and induction of the LE marker CALB1 (**Figure 5**). This stepwise switch represents a transient hybrid state in which cells co-express features of both GE and LE cells. The observed morphological switch from cuboidal GE cells to pseudostratified columnar LE cells is consistent with this interpretation. Importantly, uterine tissues obtained from control mice that lack expression of PGRMC2-3XFLAG showed similar PGR, ESR1, FOXA2 and CALB1 marker dynamics as uterine tissues from *Pgrmc2-3XFlag* mice (**Figures 4** and **5**), as well as a consistent number of junctional points (**Figure 7E**). This supports our contention that expression of the PGRMC2-3XFLAG protein does not promote or interfere with GE-to-LE transition. Despite GE-derived LE cells having the correct cellular morphology and expressing proteins consistent with developmentally-derived LE cells, their behavior and function as LE cells remains to be determined. However, we suspect that GE cells will function like LE cells once they have transitioned. In support of this, Ang *et al*. demonstrated using the *Cxcl15-iCre;R26^nTnG^*reporter mouse that following complete chemical ablation of the LE and subsequent regeneration from the GE compartment, 75% of mice were able to establish a pregnancy and maintain it to term [39].

FOXA2 has been used extensively by many labs as a marker of the GE. However, we detected FOXA2 mRNA and protein throughout the LE following postpartum endometrial regeneration (**Figure 7**). Ang *et al*. also describe expression of FOXA2 in the LE during and after endometrial regeneration [39]. FOXA2 is a highly conserved pioneer transcription factor that coordinates tissue remodeling and differentiation [55–58]. Its primary function is to bind and open condensed chromatin to expose regulatory elements that other transcriptions factors can access [59, 60]. Among its many known functions, FOXA2 regulates glucose metabolism, development of endodermal organs such as the liver and pancreas in large part by blocking epithelial-to-mesenchymal transition (EMT), and development of endometrial glands during early postnatal life [61–64]. Given its role in chromatin remodeling and blockage of EMT, it is possible that FOXA2 functions to modify chromatin structure in such a way so as to facilitate repair of the LE without it transitioning to mesenchymal tissue.

It is important to note that the existence of a *bona fide* endometrial epithelial stem cell is supported almost exclusively by *in vitro* clonal expansion studies using a standard 2D cell culture format or organoids most often consisting of single epithelial cell populations with no niche stromal support, as well as the expression of genes commonly found in other stem and non-stem cells. The expression of many of these stem cell markers, most often characterized using single cell or spatial transcriptomic platforms, changes throughout the cycle in response to sex steroids and are generally more broadly expressed rather than being uniquely expressed in a finite population of cells restricted to a unique niche as is commonly seen in other tissues that possess a stem cell population. In the absence of rigorous studies that have tested stem cell qualities of these cells *in vivo*, such as consistent identification, isolation and serial transplantation, more work is needed to identify stem cells and then determine if and how they reconstitute the endometrial epithelium during regeneration using fate mapping approaches. Sayed *et al*, provide perhaps the best support for the existence of an *Axin2^+^* progenitor cell population housed within the GE compartment that potentially gives rise to both GE and LE cells. However, this unidirectional model of propagation from a glandular progenitor to more differentiated GE and LE cells does not explain the recent finding that developmentally-derived LE cells can reconstitute the GE following conditional ablation of the entire GE with diphtheria toxin [32]. The present study neither refutes nor supports the existence of an endometrial stem cell population. An alternative to the stem cell-based theory of endometrial epithelial cell regeneration is the concept of dedifferentiation of stably, but not terminally, committed cells followed by their redifferentiation/reprogramming into cells that contribute functionally to the endometrium. As proposed in **Figure 8**, epithelial cells that are stably differentiated into GE cells to perform a specific function such as secretion of uterine histotroph during pregnancy, dedifferentiate temporarily during regeneration into uncommitted cells in response to cues produced from the underlying stromal cells and vascular tissue, as well as infiltrated immune cells recruited into the endometrium during regeneration. As the uncommitted cells integrate into the LE, they are reprogrammed to become and take on LE functions. The reverse of this process may also occur in instances where the LE is used to help reconstitute the GE during regeneration of the endometrium. In essence, GE and LE cells are maintained in semi-committed states where they perform compartment-specific functions, but have the capacity to dedifferentiate when required to facilitate endometrial regeneration depending on the needs of the tissue. This dedifferentiation/reprogramming paradigm is not unique to the endometrial epithelia. In fact, such cellular plasticity, where cells adaptively change identity, is now fully recognized as a central component of epithelial tissue renewal and repair in many organs including the esophagus, pancreas, stomach, liver, kidney and skin [65–72]. Even tissues that have a well characterized stem cell population, such as the small intestine, also use dedifferentiation/reprogramming of more committed cells as a means to rapidly repair the gut lining following excessive damage. For example, enterocytes, enteroendocrine cells, tuft cells and paneth cells all retain some level of plasticity and can switch epithelial cells fates based on need during tissue repair [73–77].

**Figure 8.**
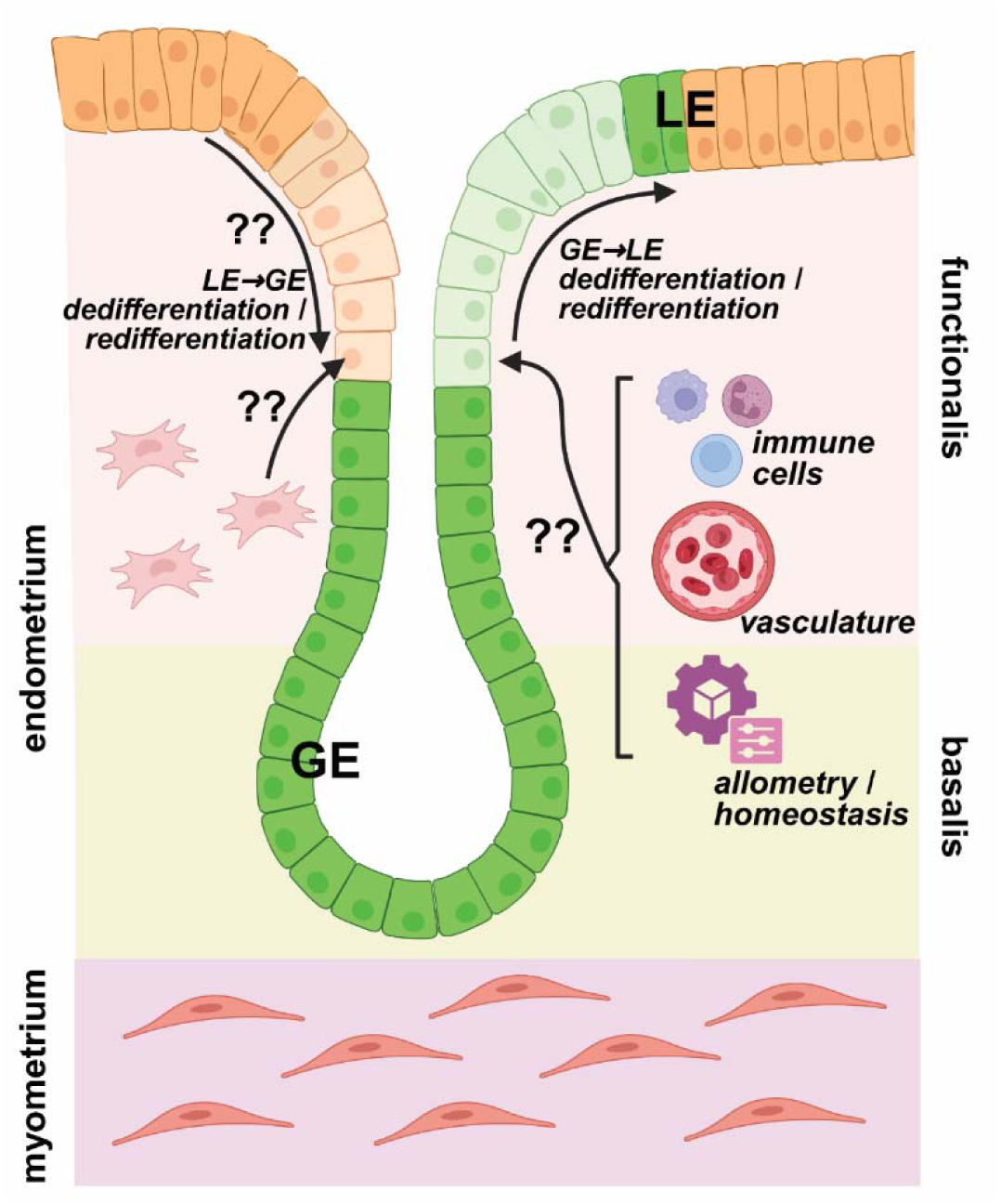
Hypothetical model of endometrial GE and LE dedifferentiation/reprogramming during postpartum regeneration. The concept that the glandular epithelium (GE) contributes to regeneration of the luminal epithelium (LE) was proposed over a century ago [33]. The schematic shows bi-directional dedifferentiation and redifferentiation of endometrial GE and LE cells during post-parturient regeneration. Using indelibly marked GE cells, the current study provides evidence that in opposition to or in conjunction with other mechanisms of endometrial regeneration such as a stem cell or bone marrow driven process, GE cells can physically contribute to the LE through processes that likely involves dedifferentiation of GE cells and their subsequent redifferentiation in LE cells. Simple cuboidal GE cells transition to pseudostratified LE cells that take on an expression profile consistent with developmentally-derived LE cells such as consistent regulation of PGR and ESR1 and CALB1. This form of epithelial regeneration is likely driven by spatial factors such as exposure to a glandular lumen versus exposure to the uterine luminal space, as well as spatial cues derived from the underlying stromal compartment that may involve stromal cell paracrine signaling molecules, or those derived from a regenerative vasculature, local immune cells and other factors that ensure maintenance of tissue allometry and homeostasis.

In conclusion, a GE-specific reporter system was used to track the fate of GE cells in the adult endometrium following single and multiple pregnancies, as well as through 5 months of age in nulliparous mice. We report that developmentally-derived GE cells have the capacity to transition into LE cells following natural damage to the endometrium. Junctional points were identified where transitioning cells expressed markers of both GE and LE cells and shape shift from cuboidal to pseudostratified columnar epithelial cells. The level of GE-to-LE transition is consistent with the extent of endometrial tissue damage. It is perhaps not surprising that transdifferentiation (*i.e.*, mesenchymal-to-epithelial transition) and dedifferentiation/reprogramming have been observed in the endometrium given the common development origin of endometrial epithelial (glandular and luminal) and stromal tissues from the mesodermal rather than epidermal germ layer. The mechanism of dedifferentiation/reprogramming offers a contrast and/or addition to other proposed mechanisms of endometrial epithelial regeneration such as a stem/progenitor cell, transdifferentiation (*i.e.*, MET), and integration of bone marrow-derived cells.

## AUTHOR CONTRIBUTION

The study was conceptualized by N.S., D.S.P., C.A.P. and J.K.P., supervised by J.K.P., written by N.S. and J.K.P., and a draft manuscript discussed and edited to final version by N.S., C.A.P., A.M.K., T.E.S., and J.K.P. Studies were conducted and analyzed by N.S., D.S.P., C.A.P., and J.K.P. The *Prss29-Cre* mouse line was provided by A.M.K., T.E.S., and J.K.P. All authors have read and agreed to the published version of the manuscript.

## ACKNOLEDGEMENTS

This study was funded in part by NIH R21HD112788, P20GM103432 and R01HD102386, USDA NIFA Functional Genomics in Animal Improvement, Food Safety and Human Health Projects 1014918, WYO-632-22, and WYO-657-24, the Curtis and Marian Rochelle Endowment in Animal Science (JKP), as well as NIH R01HD112315 (AMK) and R37HD114609 (TES).

